# The *Ghsr*^Q343X^ allele favors the storage of fat by acting on nutrient partitioning

**DOI:** 10.1101/2020.10.30.362343

**Authors:** Candice Marion, Philippe Zizzari, Raphael G.P. Denis, Rim Hassouna, Yacine Chebani, Gwenaëlle Le Pen, Florence Noble, Serge Luquet, Jacques Pantel

## Abstract

The Growth Hormone Secretagogue Receptor (GHSR) mediates key properties of the gut hormone ghrelin on metabolism and behavior. Nevertheless, most recent observations also support that the GHSR is a constitutively active G protein-coupled receptor endowed of a sophisticated tuning involving a balance of endogenous ligands. Demonstrating the feasibility of shifting GHSR canonical signaling *in vivo*, we previously reported that a model with enhanced sensitivity to ghrelin (*Ghsr*^Q343X^ mutant rats) developed fat accumulation and glucose intolerance. Herein, we investigated the contribution of energy homeostasis to the onset of this phenotype, as well as behavioral responses to feeding or pharmacological challenges, by comparing *Ghsr*^M/M^ rats to wild-type littermate rats 1) as freely behaving animals using an automated system to monitor simultaneously energy intake and expenditure, respiratory exchanges and voluntary activity and 2) in feeding and locomotor paradigms. Herein, *Ghsr*^M/M^ rats showed enhanced locomotor response to a GHSR agonist while locomotor or anorexigenic responses to amphetamine or cabergoline (dopamine receptor 2 agonist), respectively, were preserved. *Ad libitum* fed *Ghsr*^M/M^ rats consumed and conditioned for sucrose similarly to littermate control rats. In calorie-restricted conditions, *Ghsr*^M/M^ rats retained food anticipatory activity and maintained better their body weight and glycemia. Finally, prior to fat accumulation *Ghsr*^M/M^ rats showed shifted fuel preference towards carbohydrates utilization without alterations of energy intake, energy expenditure or physical activity. Overall, the present study provides proof of concept that shifted GHSR signaling can operate a specific alteration in nutrient partitioning resulting in modified balance of carbohydrate/lipid utilization.

## Introduction

The *growth hormone secretogogue receptor* (GHSR) (Howard, et al. 1996) holds a unique interest as the target of the gut hormone ghrelin (Kojima, et al. 1999), a hormone with key pharmacological properties such as GH release (Kojima et al. 1999), enhanced fat storage and enhanced food intake, mediated by its action on brain energy homeostasis centers (Nakazato, et al. 2001; Theander-Carrillo, et al. 2006; Tschop, et al. 2000) and hedonic circuits (Abizaid, et al. 2006; Jerlhag, et al. 2007; Naleid, et al. 2005). While the ablation of the *Ghsr* gene (*Ghsr*^-/-^) or of ghrelin-producing cells in mice reported mitigated results on energy homeostasis (Muller, et al. 2015), *Ghsr*^-/-^ mice failed to enhance locomotor activity similarly to control mice during scheduled feeding (Blum, et al. 2009; LeSauter, et al. 2009) and to preserve glycemic control during severe caloric restriction (Wang, et al. 2014). The GHSR could therefore play key roles in situations of stress while its role in the fed state remains largely unknown (Mani and Zigman 2017).

The GHSR is a G protein-coupled receptor (GPCR) showing high constitutive activity as documented in cellular (Holst, et al. 2003) and acellular systems (Damian, et al. 2012). Recent observations identified the liver hormone LEAP2 as a novel GHSR ligand (Ge, et al. 2018) with inverse agonist properties (M’Kadmi, et al. 2019; Mani, et al. 2019). Therefore, according to its expression pattern in the periphery and in several key brain regions, including the hypothalamus, ventral tegmental area (VTA) or hippocampus (Zigman, et al. 2006), the GHSR could exert its functions according to a complex balance between GHSR ligands, whose accessibility in each GHSR expressing brain structure still needs to be delineated (Perello, et al. 2019). Overall, these observations, that appear as a game changer (Andrews 2019), support an unprecedented refinement for an endocrine system, questioning the therapeutic potential of this GPCR target (Al-Massadi, et al. 2018). We reasoned that a preclinical model with shifted canonical GHSR signaling could provide key insights for future drug discovery.

To this purpose, we used *Ghsr*^Q343X^ mutant rat model in which GHSR receptor display shifted signaling towards G protein pathways characterized notably by increased GH release and chow intake in response to GHSR agonists. Homozygous mutant rats carrying this allele (*Ghsr*^M/M^) develop fat accumulation and insulin resistance with age, a phenotype consistent with enhanced ghrelin effects (Chebani et al. 2016). However, the contribution of energy homeostasis to the onset of fat accumulation in *Ghsr*^M/M^ rats remained unexplored, as well as the physiological consequences of this mutation on food-related behaviors. In this study, we assessed the consequence of shifting GHSR signaling onto 1) feeding and locomotor response to pharmacological manipulations or feeding challenges, and 2) metabolic efficiency, nutrient partitioning in automated food and calorimetric recording system.

## Material and methods

### Animals

The mutant line of FHH-Ghsr^m1Mcwi^ rats carrying the *Ghsr*^Q343X^ allele was obtained from the Medical College of Wisconsin (Milwaukee, WI, USA). The homozygous FHH-*Ghsr*^m1Mcwi^ rats and wild-type littermates are referred to as *Ghsr*^M/M^ and *Ghsr*^WT/WT^ rats, respectively. Animals used in this study (187 rats from 7 litters) were obtained from crossing heterozygous rats. Animals were raised four by cages with free access to water and chow diet (A04, SAFE), in a room with controlled temperature (22-24°C) and illumination (12h light/dark schedule with lights on at 7:00 am). The genotype of the rats at the *Ghsr* locus was determined as previously described (Chebani et al. 2016). Experiments were performed with male or female rats with *ad libitum* access to food and water, unless otherwise specified. The procedures involving rats were approved by the ethics committee on animal experimentation of the Université Paris-Descartes.

### Amphetamine-induced locomotion

A group of experimentally naive 10 week-old male rats was used to assess the locomotor response to a novel environment and to the injection of the dopamine enhancing drug amphetamine. Rats were habituated to a locomotor cage (Imetronic, Bordeaux, FR) for 60 mins on 2 consecutive days. The next 2 days, groups of *Ghsr*^M/M^ and *Ghsr*^WT/WT^ rats were injected i.p. with amphetamine (2 mg/kg) while a control group of *Ghsr* heterozygous rats was injected i.p. with saline.

### Cabergoline-induced anorexigenic response

Prior to the food intake experiment, 7 month-old female *Ghsr*^M/M^ and *Ghsr*^WT/WT^ rats were habituated to single housing for at least 1 week. Free access to chow diet was removed at 6:00 pm for a 16h fast. Rats were then s.c. injected with the dopamine receptor D2 (DRD2) agonist cabergoline (0.5 mg/kg, Tocris) or saline at 11:00 am. Each animal received both treatments in a crossover fashion, with a 2-week washout period between treatments.

### Hexarelin-induced locomotion

On the first day of each experiment, experimentally naive 5 month-old male rats were individually put in an open field arena (Med Associates Inc., Vermont, USA) with clean bedding. Room illumination was set at 15 lux. Recording sessions began each day at 8:30 am. Rats were habituated to the apparatus on 2 consecutive days for 45 min sessions. The next day, rats were injected by injected s.c. with saline solution (100 μL/100mg). Every other day, rats were challenged with varying doses of hexarelin (10 – 300 nmol/kg, Sigma) in the same setup, using a random crossover design.

### Blood ghrelin measurements

Blood samples were collected from the same cohort of rats in a second experiment performed 3 weeks after the calorimetry exploration. Blood samples were obtained by tail bleeding in *ad libitum* fed, 24h fasted and 24h refed conditions. Sample preparation and ghrelin measurements were performed as previously reported (Chebani et al. 2016).

### Sucrose two-bottle choice

*Ad libitum* fed 7 month-old male rats were habituated to single housing and drinking from 2 bottles of water for at least 1 week prior to the beginning of the experiment. On 7 consecutive days they had access to a 0.75% sucrose solution and water for 1h, starting at 10:00 am. The position of the sucrose bottle was switched every day to prevent habituation. Bottles were weighed before presentation and again at the end of the 1h test to assess fluid intakes.

### Operant responding for sucrose pellets

Operant chambers (Med Associates Inc., Vermont, USA) comprised 2 open nose-pocking niches, one designed as the active target (triggering reward delivery) and the other as the inactive, dummy target. Entry into the active nose-poking hole was rewarded by the delivery of a 45 mg sucrose pellet (Dustless Precision Pellets^®^). Entries into the inactive nose-poking hole had no consequence. House-light was on throughout the session. A light was switched on in the active nose-poking hole or food magazine to cue target or reward availability, respectively.

*Ad libitum* fed 7 month-old female rats were trained on fixed ratio (FR) schedules, starting with FR1 (1 nose-poke required to obtain 1 pellet), then on FR3 (3 nose-pokes to obtain 1 pellet) and finally FR5. A training session ended after 30 mins or when the rat had obtained 50 pellets (success criterion). Rats were kept on each FR schedule for 4 consecutive sessions, at the end of which all animals had reached the success criterion. After training, the rats underwent consecutive test sessions on a progressive ratio (PR) schedule to assess their motivation to work for the food reward according to the usual exponential formula (N = 5 * exp(0.2N) – 5). The session ended when rats had failed to obtain a pellet for 30 consecutive minutes.

### Running wheels

7 month-old male *Ghsr*^WT/WT^ and *Ghsr*^M/M^ rats matched for body weight were individualized at least 1 week prior to the experiment. Body weight, chow intake and wheel running behavior were monitored daily for 13 days under *ad libitum* feeding conditions until daily running levels had stabilized. On the last baseline day, chow intake was removed at 1:00 pm, and rats were fasted for 24h. On the following days, rats had free access to chow 2h/day, from 1:00 to 3:00 pm. During food access, running wheels were blocked to make sure the rats would feed and not run. The rats had free access to the running wheel for the 22 remaining hours. Every quarter of wheel turn was detected and recorded using a Matlab program. Raw data were converted to numbers of wheel turns in 1h bins using a dedicated R program. Chronic food restriction lasted for 15 days before rats were euthanized.

### Food anticipatory activity assessed in open field arena

7 month-old male rats were individualized at least 1 week prior to the experiment. On the first session, locomotor activity was recorded in the open field arena for 2 hours before rats were returned to their homecage where food had been removed. During the 20 following days, rats were recorded daily in the same arena for 2h (bedding left in place) before being returned to their homecage and having access to chow diet for 4h. Body weight and chow intake were assessed daily throughout the experiment.

### Analysis of Metabolic efficiency

Calorimetry exploration was performed using 10 week-old male rats at the start of the experiment. Body composition was assessed at the start of the experiment, at the end of baseline, after 24h fasting, and after 48h refeeding (end of the experiment) using an Echo Medical system (EchoMRI 100, Whole Body Composition Analyser, EchoMRI, Houston, USA). Energy expenditure, oxygen consumption and carbon dioxide production, respiratory exchange ratio, food intake and homecage activity were obtained using calorimetric chambers (Labmaster, TSE Systems GmbH, Bad Homburg, Germany) as previously depicted (Joly-Amado, et al. 2012). Rats were individually housed, fed standard chow and acclimated to the chambers for 48h before experimental measurements. A meal was defined as the consumption of > 0.3 g of food, separated from the next feeding episode by at least 10 mins. To assess metabolic flexibility, all RER data were compiled to obtain relative cumulative frequency curves for *Ghsr*^M/M^ and *Ghsr*^WT/WT^ rats. Sigmoidal dose-response curves were fitted to determine EC_50_ and Hill slopes (indicative of metabolic rigidity) (Riachi, et al. 2004).

### Statistics

Results are presented as mean ± SEM. Sample size (n) and *p* values are given in the figure captions. Statistical analyses were performed using GraphPad Prism^®^ 5.01, SPSS statistics (IBM Inc.) and R software. Differences between 2 groups were determined using non-parametric Mann-Whitney or Wilcoxon tests, or parametric Student’s tests, as appropriate. Comparisons of multiple groups were performed using two-way ANOVA, repeated measure ANOVA or ANOVA on aligned rank transformed data (ARTool package) and *p* value of post-hoc tests were adjusted with the Sidak correction. Covariance analyses (ANCOVA) used the Mouse Metabolic Phenotyping Center web page http://www.mmpc.org/shared/regression.aspx). For all statistical analyses, a *p* value of less than 0.05 was considered as significant.

## Results

### Pharmacological probing of central dopaminergic circuits in *Ghsr*^M/M^ rats reveals enhanced locomotor response to a GHSR agonist but preserved responses to amphetamine or to a DRD2 agonist

To examine central dopaminergic circuits in *Ghsr*^M/M^ rats, locomotion or refeeding response was examined in *Ghsr*^M/M^ and *Ghsr*^WT/WT^ rats in response to pharmacologic modulators of dopamine signaling or GHSR agonist. First, we assessed the locomotor response to amphetamine (AMPH), a psychoactive drug known to reverse dopamine transporter activity at post-synaptic target leading to enhances DA release and action (Fig. 1A, B, C). Both experimental groups displayed similar response to the novel environment (Fig. 1A), and to AMPH-induced hyperlocomotion (Fig. 1B, C). Second, GHSR-DRD2 heteromers can produce behavioral response independent of ghrelin binding and correlated to DA function (Kern, et al. 2012). *Ghsr^M/M^* rats model offer a great platform to probe how possible interaction of GHSR and DRD2 receptor might contribute to alter response to DRD2 agonist.To this purpose, the anorexigenic effect of the DRD2 agonist cabergoline was evaluated in *Ghsr*^M/M^ and *Ghsr*^WT/WT^ rats. Cabergoline mediated a potent anorexigenic response in the refeeding of prefasted *Ghsr*^M/M^ and *Ghsr*^WT/WT^ animals (Fig. 1D, E). No significant difference were found across genotypes. Third, the pharmacological properties of the GHSR on locomotion were tested in *Ghsr*^M/M^ and *Ghsr*^WT/WT^ rats challenged with varying doses of the agonist hexarelin. Repeated measure ANOVA revealed a significant dose x genotype interaction (p<0.01) suggestive of a differential dose-response effect between the two groups of rats (Fig. 1F). Indeed, post hoc analyses showed that the locomotor response to the highest tested dose of hexarelin was twice as high in *Ghsr*^M/M^ compared to *Ghsr*^WT/WT^ littermate rats, an observation further confirmed in the analyses of the response to the highest hexarelin dose (Fig. 1G, H). Overall, these observations, while supporting enhanced GHSR responsiveness in *Ghsr*^M/M^ rats, also rule out gross abnormalities in the dopaminergic system of these rats.

**Figure 1:**
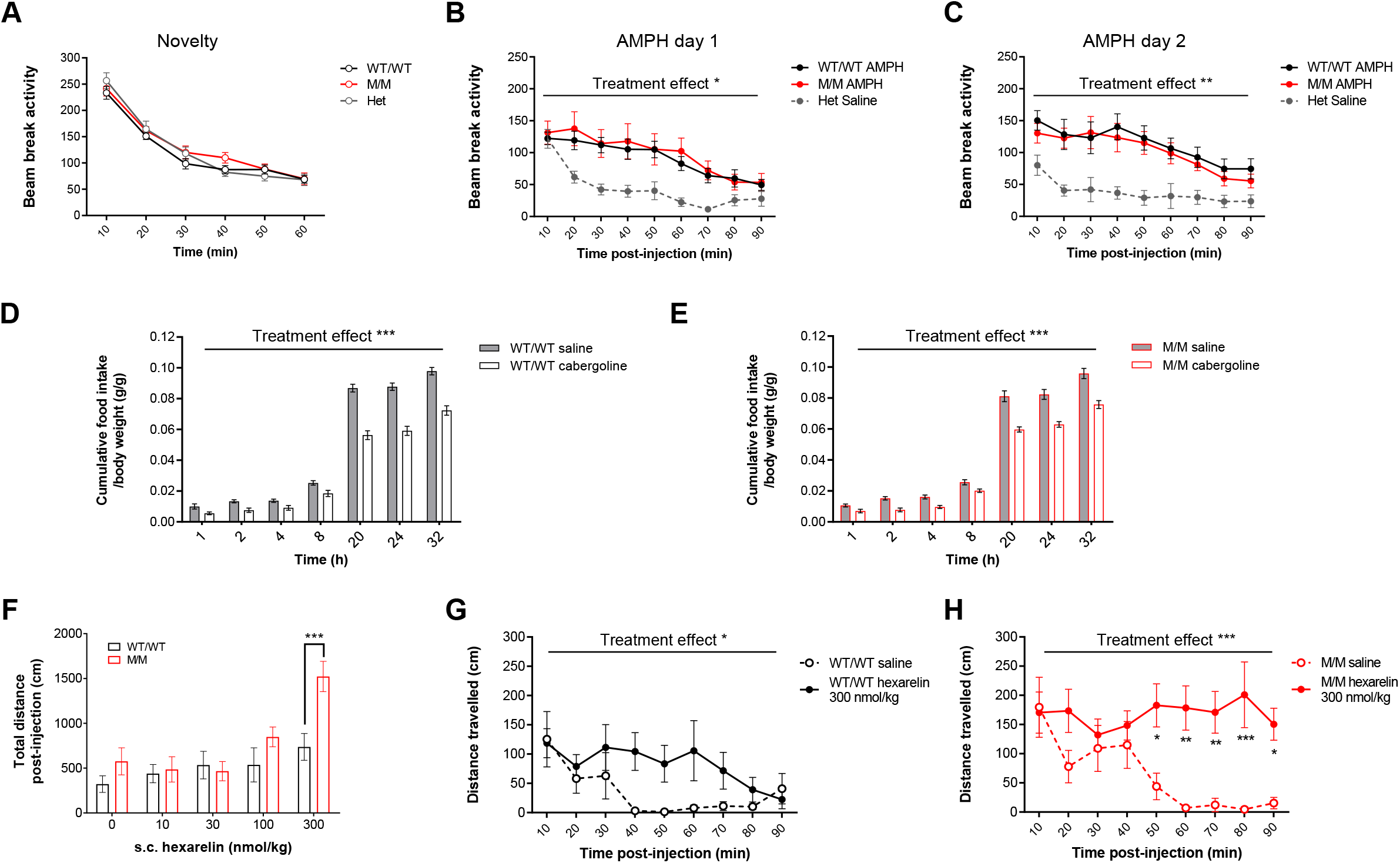
Pharmacological probing of central dopaminergic circuits in *Ghsr*^M/M^ rats. (A) Time course of the locomotor response to a novel environment and (B-C) to the injection of i.p. amphetamine (2 mg/kg) on two consecutive days in *Ghsr*^M/M^ (n=11) and *Ghsr*^WT/WT^ rats (n=10), *Ghsr* heterozygous rats served as the saline control group (n=7). Anorexigenic effects of a DRD2 agonist in a refeeding paradigm performed in fasted *Ghsr*^WT/WT^ (n=9) (D) and *Ghsr*^M/M^ (n=9) (E) rats injected s.c. with cabergoline (0.5 mg/kg) or with saline before refeeding. (F) Total locomotor response to increasing doses of the GHSR agonist hexarelin s.c. injected in male *Ghsr*^WT/WT^ (n=13) and *Ghsr*^M/M^ (n=13) rats, and (G-H) locomotor responses obtained with the highest injected hexarelin dose or saline across time. Data were analyzed by 2-way repeated measure ANOVA followed by Sidak’s post-hoc tests. * p<0.05; ** p<0.01; *** p<0.001; ~ non-significant trend (p<0.1). Data represent mean ± SEM.

### *Ghsr*^M/M^ rats show unaltered spontaneous conditioning and motivation for sucrose but accelerated performance in operant system

The former results indicated that relative to their body weight, chow consumption was comparable between *Ghsr*^M/M^ and *Ghsr*^WT/WT^ rat, while showing enhanced sensitivity to endogenous ghrelin (Chebani et al. 2016). We hypothesized that these animals might show improved consumption and/or motivation for palatable food. To test this, we took advantage of the spontaneously high preference for sucrose in the fawn hood strain (Tordoff, et al. 2008) to investigate the spontaneous consumption and reinforcing properties of sucrose in *Ghsr*^M/M^ and *Ghsr*^WT/WT^ rats using free choice and instrumental conditioning paradigms. In a two-bottle choice between a sucrose solution and water, both *Ghsr*^M/M^ and *Ghsr*^WT/WT^ male littermates displayed similar levels of consumption throughout days, with a strong, sustained preference for the sucrose solution over water (Fig. 2A). For instrumental conditioning paradigms, we focused our interest on female rats who successfully achieved conditioning without requiring calorie restriction, a manipulation known to interfere with ghrelin endogenous tone. During conditioning sessions, *ad libitum* fed *Ghsr*^M/M^ and *Ghsr*^WT/WT^ rats achieved success criterion during the last 2 sessions of each Fixed Ratio (FR) schedule (Fig. 2B upper panel; repeated measure ANOVA; session effect; p<0.001). Total responding levels were similar between *Ghsr*^M/M^ and *Ghsr*^WT/WT^ rats on the Progressive Ratio (PR) PR schedule, and for both groups, nose-poking activity on the inactive, dummy target was close to zero, indicating specific, goal-oriented responding at the active nose-poking hole. Interestingly, at 5 mins after the beginning of sessions, *Ghsr*^M/M^ rats had significantly achieved more responses than *Ghsr*^WT/WT^ rats, starting from the 3^rd^ FR3 session (Fig. 2B lower panel), although the total number of responses was comparable between *Ghsr*^M/M^ and *Ghsr*^WT/WT^ rats throughout the FR and PR schedules. This significant difference in performing speed was also seen 15 mins after the beginning of the session (Fig. 2B lower panel). Overall, *ad libitum* fed *Ghsr*^M/M^ rats showed preference or consumption for palatable food similar to *Ghsr*^WT/WT^ rats, but performed faster to obtain rewarding food in an operant nose-poking task.

**Figure 2:**
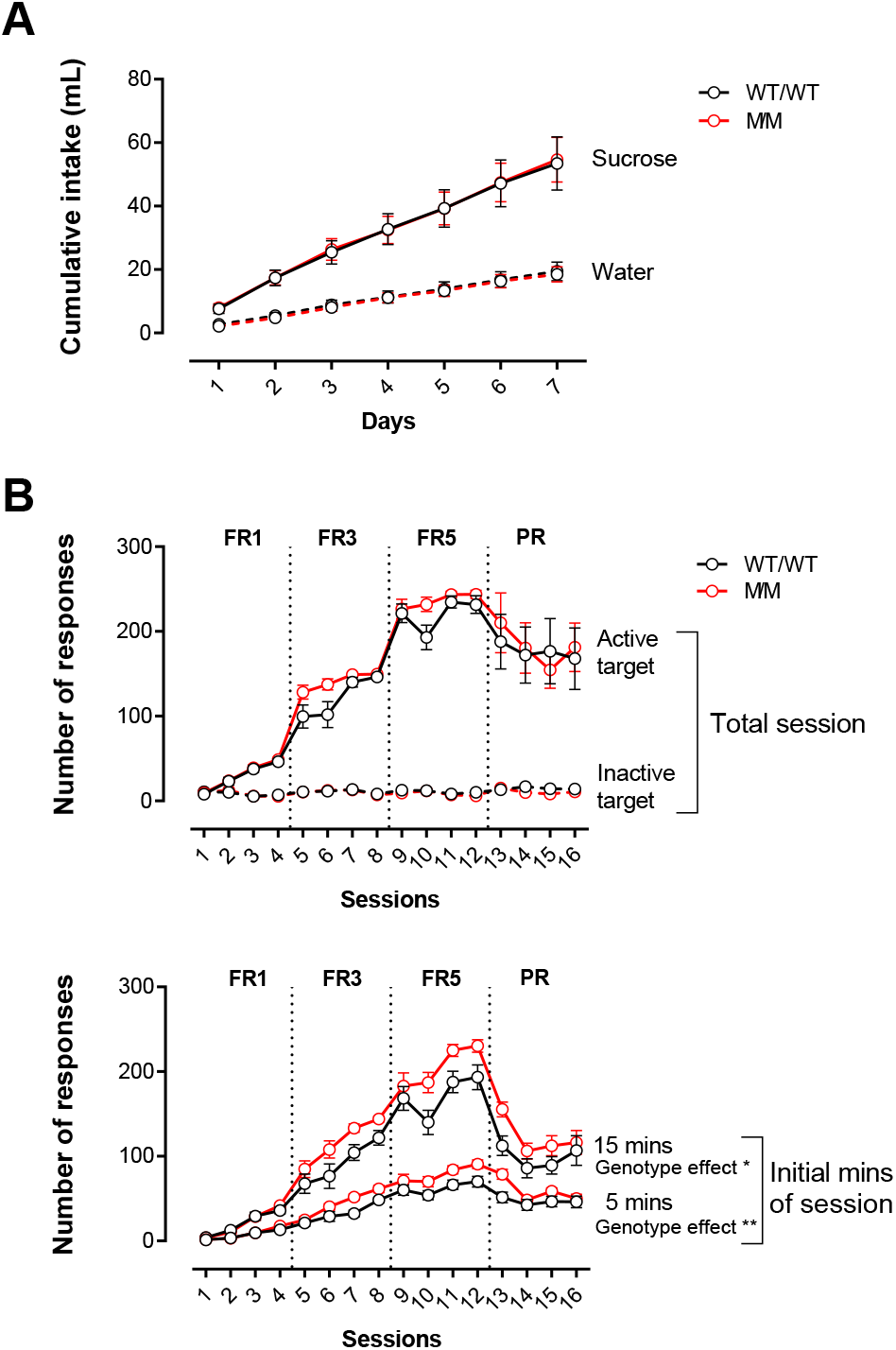
Consumption and motivation for sucrose in *Ghsr*^M/M^ rats. (A) Cumulative intake of sucrose solution (0.75%) (plain lines) and water (dotted line) during daily 1h two-bottle choice in *Ghsr*^M/M^ (n=17) and *Ghsr*^WT/WT^ (n=13) *ad libitum* fed rats. (B) Operant nose-poking responses for sucrose pellets in *Ghsr*^M/M^ (n=16) and *Ghsr*^WT/WT^ (n=15) *ad libitum* fed rats during each session (upper panel) or at intermediate times of the session (lower panel). Data were analyzed by 2-way repeated measure ANOVA.* p<0.05; ** p<0.01. Data represent mean ± SEM.

### *Ghsr*^M/M^ display preserved food anticipatory activity in response to scheduled feeding

Previous investigations revealed that *Ghsr*^M/M^ rats preserved better their body weight and glycemia in scheduled restricted feeding conditions (Chebani et al. 2016), however whether this condition also affected other behavioral response related to food-seeking response remained unexplored. To test this, two paradigms differing on the kind of locomotor activity measured (wheel running or ambulatory locomotion) were used. In the first experiment, *Ghsr*^M/M^ and *Ghsr*^WT/WT^ littermate rats habituated to a single housing cage with free access to running wheels were put on a restricted feeding schedule. Unexpectedly, during the *ad libitum* feeding period, *Ghsr*^M/M^ rats showed decreased running activity during the first hour of dark phase (Fig. 3A). At the beginning of caloric restriction, wheel running activity was increased in both groups of rats and the activity of *Ghsr*^M/M^ rats no longer differed from that of *Ghsr*^WT/WT^ rats (Fig. 3B). Running activity was further enhanced at the end of the caloric restriction and the activity pattern was re-distributed with maximal activity levels in anticipation to dark phase and meal access (Fig. 3C). *Ghsr*^M/M^ and *Ghsr*^WT/WT^ rats displayed comparable levels of daily activity, as well as similar food anticipatory behavior (Fig. 3C). Besides, throughout caloric restriction *Ghsr*^M/M^ rats consumed similar amounts of food compared to *Ghsr*^WT/WT^ littermates (Fig. 3D) but were better able to maintain their body weight (Fig. 3E) and glycemia (Fig. 3F). To verify that food anticipatory activity was indeed unaltered in *Ghsr*^M/M^ rats while avoiding the potentially confounding effect of wheel running, which is rewarding by itself, we used an alternative paradigm in which another cohort of rats at same age were put on a 4h restricted feeding schedule and recorded daily in actimetry cages during the 2 hours preceding food access. In this setting, *Ghsr*^M/M^ and *Ghsr*^WT/WT^ rats developed food anticipatory activity to comparable levels (Fig. 3G), while chow intake and body weight were similar across genotypes (Fig. 3H, I). Altogether, *Ghsr*^M/M^ rats therefore show anticipatory activity and food consumption similar to *Ghsr*^WT/WT^ littermates on a restricted feeding schedule, suggesting that the *Ghsr*^Q343X^ mutation does not affect behavioral anticipation of food.

**Figure 3:**
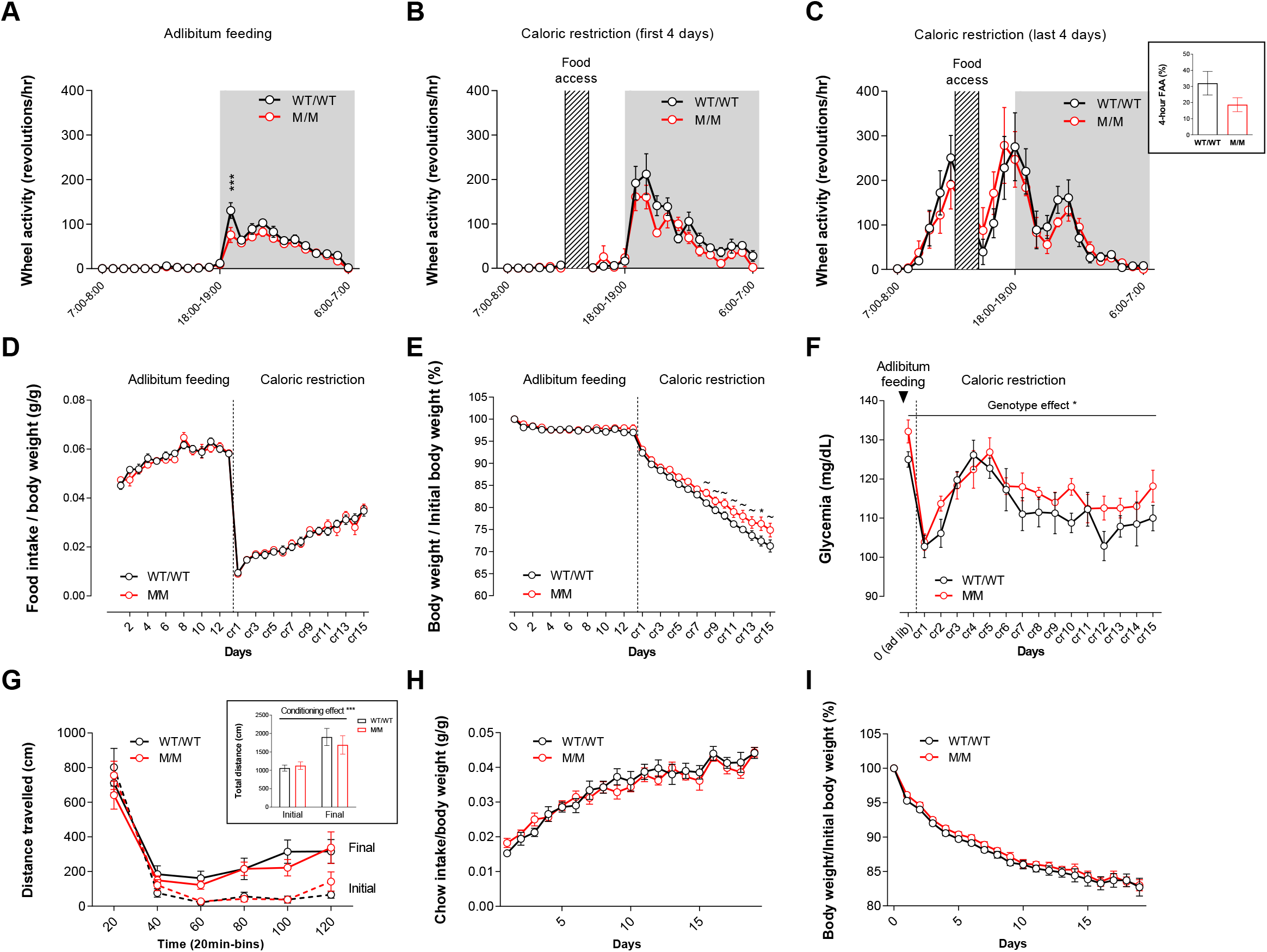
Food anticipatory activity is preserved in *Ghsr*^M/M^ rats. Wheel running activity in *Ghsr*^M/M^ (n=7) and *Ghsr*^WT/WT^ (n=8) rats averaged over 10 days of *ad libitum* feeding (A) or at beginning (B) or end of the restricted feeding schedule (C), daily food intake (D), body weight (E) and glycemia (F) of rats across the protocol. Travelled distance recorded in an open field in the 2 hours preceding food access in *Ghsr^M/M^* (n=8) and *Ghsr*^WT/WT^ (n=7) rats put on a 4h restricted feeding schedule (initial: averaged on the first 4 days; final: averaged on the last 4 days) (G), daily food intake normalized to body weight (H) and body weight (I). Data were analyzed by 2-way repeated measure ANOVA followed by Sidak’s post-hoc tests. * p<0.05; ** p<0.01; *** p<0.001; ~ non-significant trend (p<0.1). Data represent mean ± SEM.

### *Ghsr*^M/M^ rats display increased body mass and similar body composition at 12 weeks of age

In order to determine whether alteration in energy homeostasis precede the development of insulin resistance in 6 month-old rats (Chebani et al. 2016), metabolic efficiency was assessed using indirect calorimetry in asymptomatic mutant rats in response to nutritional manipulation. *Ghsr*^M/M^ and *Ghsr*^WT/WT^ rats entered in metabolic cages at similar age (12.3±0.2 and 12.0±0.3 week-old, respectively). Upon entry, *Ghsr*^M/M^ rats were on average 7% heavier than *Ghsr*^WT/WT^ littermates, but proportions of fat and lean masses relatively to total body mass were comparable in *Ghsr*^M/M^ and *Ghsr*^WT/WT^ rats (Fig. 4A). Across the experiment, *Ghsr*^M/M^ and *Ghsr*^WT/WT^ animals similarly lost and regained total body mass, fat mass and lean mass upon fasting and refeeding, respectively (Fig. 4B). Thus, in young adult rats, the *Ghsr*^Q343X^ mutation resulted in higher body weight, with increases in both fat and lean masses in proportion to total body mass, suggesting an aggravation of the phenotype later with age, which is in line with similar blood glucose levels between genotypes in all feeding conditions (Supplementary Fig. 1). As shown Fig. 4C, *Ghsr*^M/M^ and *Ghsr*^WT/WT^ animals showed comparable 24h patterns of activity across baseline, fasting and refeeding, both in light and dark phases. In both groups of rats, fasting increased homecage activity, while refeeding returned it to basal level (Fig. 4D). As expected, the feeding pattern post-fast significantly differed from the *ad libitum* pre-fast condition for both groups of rats (Fig. 4E), with no differences across genotypes regarding 24h intake (Fig. 4F). Furthermore, the diurnal and nocturnal meal parameters across feeding conditions were comparable between *Ghsr*^M/M^ and *Ghsr*^WT/WT^ rats, as measured by mean meal size, total time spent eating, number of meals, meal duration, inter-meal intervals and ingestion rate (Supplementary Fig. 2). Not surprisingly, both groups of rats showed similar diurnal and nocturnal levels of energy expenditure characterized by decreased energy expenditure during fasting, while there was a general tendency to return to its basal level upon refeeding (Fig. 4G). Analyses of 24h energy expenditure revealed no differences between genotypes in the *ad libitum* feeding condition, but a trend to decreased energy expenditure was observed in *Ghsr*^M/M^ rats during fast and refeeding conditions (Fig. 4H). Estimated resting metabolic rate was also comparable for *Ghsr*^M/M^ rats and their *Ghsr*^WT/WT^ littermates (9.1±0.4 kcal/h/lean mass and 9.4±0.3 kcal/h/lean mass, respectively). Altogether, these results suggest that in young adult rats, the *Ghsr*^Q343X^ mutation does not alter daily locomotor activity, caloric intake, meal patterns or energy expenditure in conditions of *ad libitum* access to food as well as short-term food deprivation.

**Figure 4:**
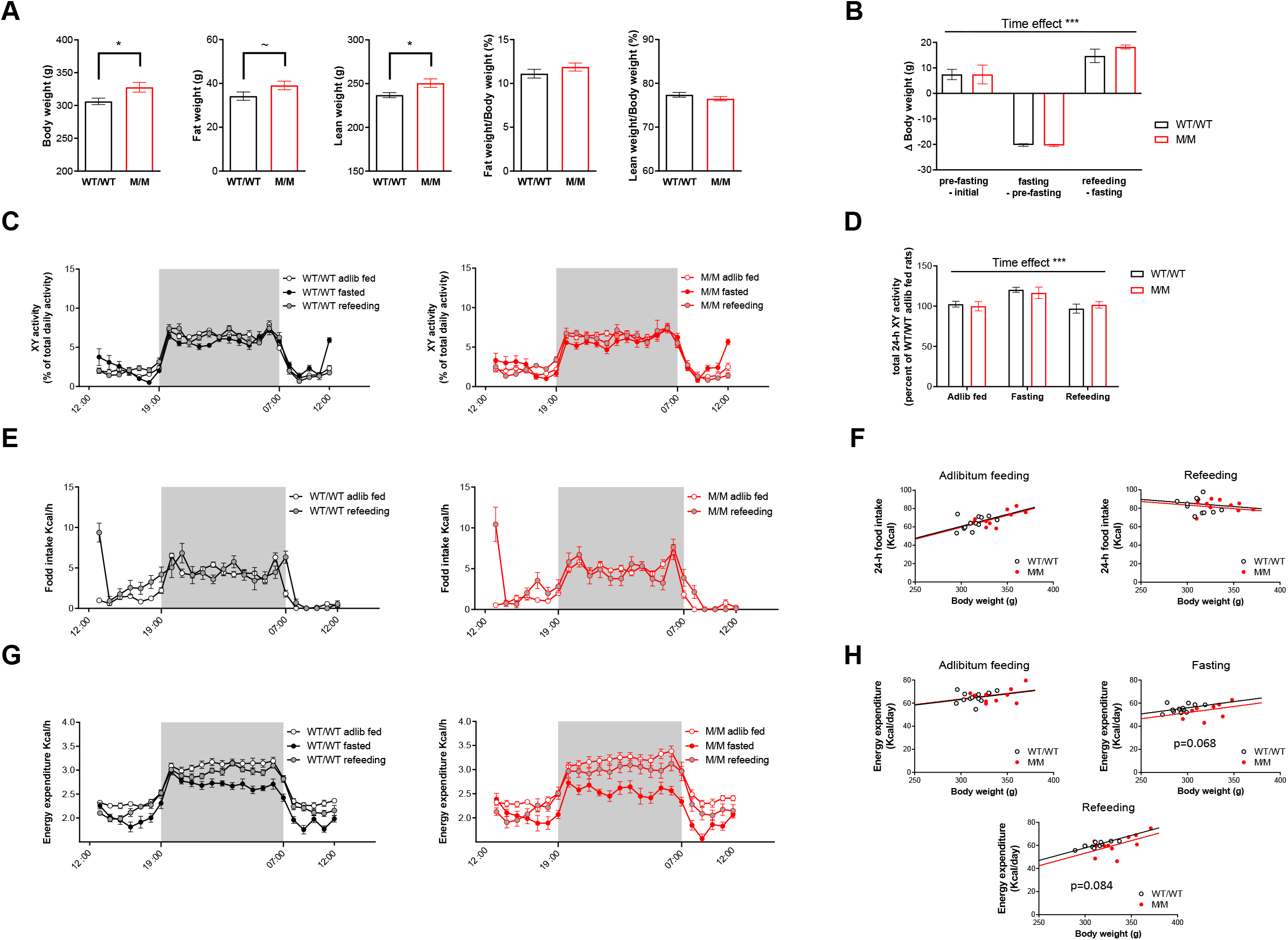
*Ghsr*^M/M^ rats show preserved circadian locomotor, feeding and energy balance rhythms across feeding conditions. (A) Body composition at the start of the calorimetric experiment in young adult (12 week-old) rats (n=12/genotype). (B) Changes in total body weight across *ad libitum* feeding (4 days), fasting (24h) and refeeding (48h). (C) Daily fluctuations of homecage activity and (D) 24h homecage activity. (E) Daily fluctuations of food intake and (F) 24h food intake as a function of body mass in each rat. (G) Daily fluctuations of energy expenditure and (H) 24h energy expenditure as a function of body mass in each rat. Values represent means ± SEM. Data were analyzed by Mann-Whitney test (A), by ANOVA on aligned rank transformed data (B-E; G) or by ANCOVA using body weight as a covariate (F; H). * p<0.05; *** p<0.001; ~ non-significant trend (p<0.1). Data represent mean ± SEM.

### *Ghsr*^M/M^ rats exhibit metabolic fuel preference towards carbohydrates

As *Ghsr*^M/M^ rats showed increased body weight without any major changes in energy intake or expenditure compared to their *Ghsr*^WT/WT^ littermates, we then investigated if *Ghsr*^M/M^ and *Ghsr*^WT/WT^ rats displayed differences in utilization of metabolic substrates, by exploring respiratory exchange ratio (RER) in baseline, fasted and refed conditions (Fig. 5A, B, C). In all nutritional states, mean 24h RER was higher in *Ghsr*^M/M^ rats compared to *Ghsr*^WT/WT^ littermates (Fig. 5D), indicating a slight but sustained increase in carbohydrate utilization as energy substrate, at the expense of fat, in *Ghsr*^M/M^ rats compared to *Ghsr*^WT/WT^ rats. However, qualitative RER variations across the experiment were similar in *Ghsr*^M/M^ and *Ghsr*^WT/WT^ animals, with RER decreasing during fasting compared to *ad libitum* basal feeding (Fig. 5A, B, D), and returning to baseline levels during the first 24h of refeeding (Fig. 5C, D). Separate analyses of light and dark phases indicated that, during light phase, *Ghsr*^M/M^ rats had in all conditions a higher RER than *Ghsr*^WT/WT^ rats (~2% in the fed state) (Fig. 5E) which is not the case during the dark phase (Fig. 5F). Thus, *Ghsr*^M/M^ compared to *Ghsr*^WT/WT^ rats presented an overall decreased use of lipids and increased use of carbohydrates as energy substrates illustrated further by decreased cumulative fat oxidation across feeding conditions (Supplementary Fig. 3). In order to explore metabolic flexibility, RER data were compiled to obtain relative cumulative frequency curves for *Ghsr*^M/M^ and *Ghsr*^WT/WT^ rats. Comparison of the fits revealed a right shift of the curve for *Ghsr*^M/M^ compared to *Ghsr*^WT/WT^ rats (Fig. 5G), indicating that RER distribution of the *Ghsr*^M/M^ group is narrower and skewed towards higher values compared to the *Ghsr*^WT/WT^ group. Moreover, 1/Hill slope was lower for *Ghsr*^M/M^ rats compared with *Ghsr*^WT/WT^ rats indicative of decreased metabolic flexibility in *Ghsr*^M/M^ rats. To complete this study, blood concentrations of acyl and des-acyl ghrelin isoforms were assessed using the same animals in satiated, fasted and refed states (Fig. 5H, I). *Ghsr*^M/M^ and *Ghsr*^WT/WT^ littermates showed similar blood acyl and des-acyl ghrelin concentrations before fasting and after 24h of refeeding. In contrast, the fasting-induced increase in both ghrelin isoforms was lower in *Ghsr*^M/M^ rats compared to *Ghsr*^WT/WT^ rats, while acyl ghrelin to total ghrelin ratios remained comparable in both groups of rats. Altogether, phenotypic observations obtained using the automated system in *Ghsr*^M/M^ rats are likely to occur at similar or even decreased circulating ghrelin levels in fed or fasted rats, respectively.

**Figure 5:**
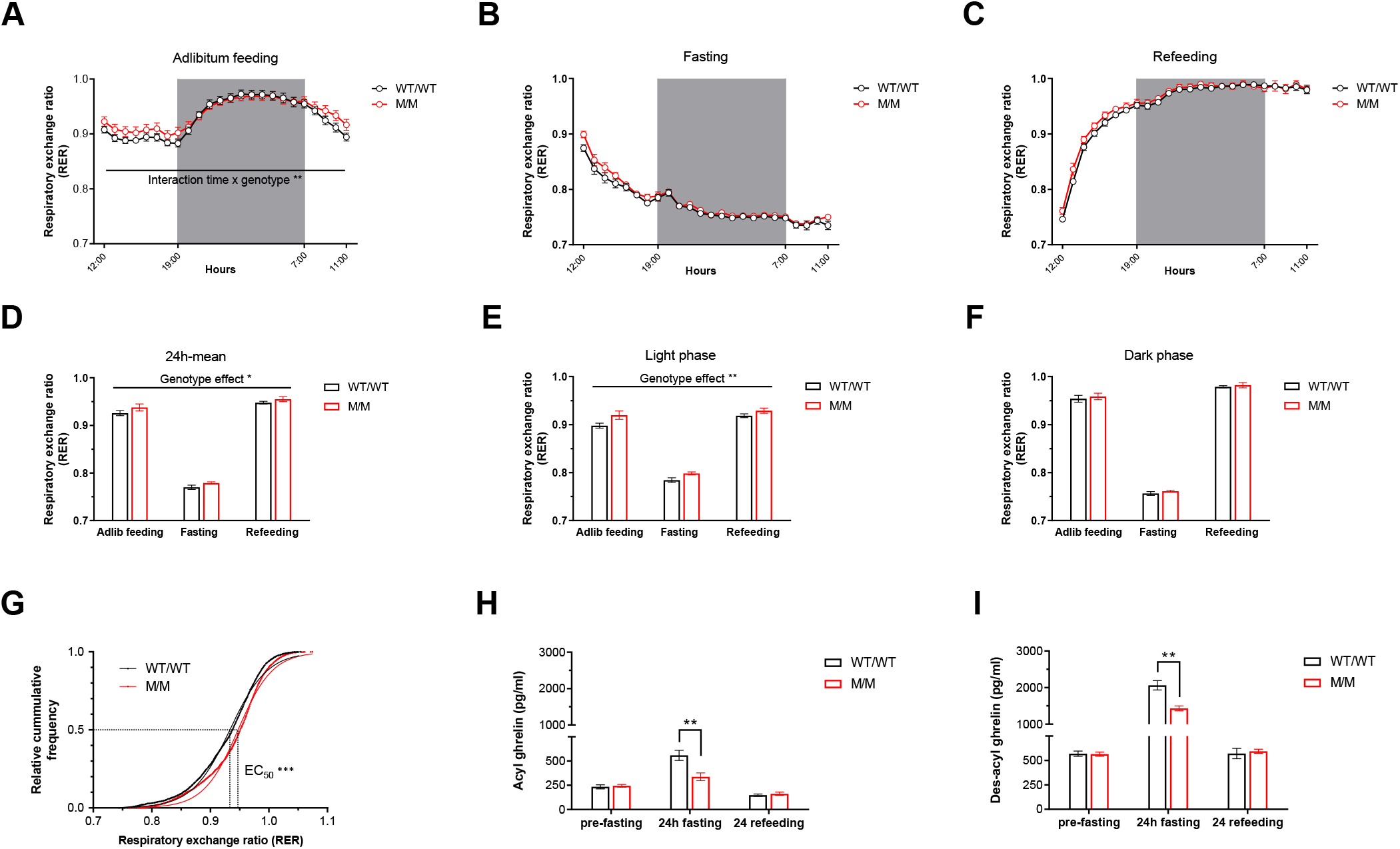
Increased respiratory exchange ratio in *Ghsr*^M/M^ rats across feeding conditions. Daily pattern of respiratory exchange ratio (RER) during adlibitum condition (4-day average) (A), 24h fasting (B) and the first 24h of refeeding (C) in 12 week-old male rats (n=12/genotype). Averaged RER over 24h (D), light phase (E) and dark phase (F) for each feeding condition. (G) Relative cumulative frequency curves for *ad libitum* fed *Ghsr*^M/M^ and *Ghsr*^WT/WT^ rats (dots) and curve fits (full lines) reveal higher EC_50_ (0.947 vs. 0.933) and a lower 1/Hill slope value (0.067 vs. 0.074) for *Ghsr*^M/M^ rats compared to *Ghsr*^WT/WT^ rats (F-test; p<0.001 for both parameters). (H-I) Blood ghrelin isoforms during ad libitum feeding, fasting and refeeding in the same rats. Data were analyzed by ANOVA on aligned rank transformed data (A-F) or 2-way ANOVA (H,I). * p<0.05; ** p<0.01; *** p<0.001. Data represent mean ± SEM.

## Discussion

Recent observations suggest that the GHSR is a constitutively active GPCR endowed of a sophisticated tuning involving by a balance of endogenous ligands (Ge et al. 2018; Mani et al. 2019). Therefore, shifting GHSR canonical signaling could provide unique insights for future drug discovery. This is the case of the functionally significant *Ghsr*^Q343X^ mutation, whose homozygous rat carriers display enhanced responsiveness to GHSR agonists, enhanced fat accumulation, and insulin resistance (Chebani et al. 2016). The present study now shows that *Ghsr*^M/M^ rats specifically display an enhanced locomotor response to a GHSR agonist while responses using dopaminergic drugs are seemingly unaltered. Similarly, spontaneous consumption and conditioning for sucrose appeared not to be impacted by the *Ghsr*^Q343X^ mutation, nor did food anticipatory activity. *Ghsr*^M/M^ rats, prior to fat accumulation and insulin resistance, show a shifted fuel preference towards carbohydrates at early adulthood. In contrast, *Ghsr*^M/M^ rats did not show obvious qualitative or quantitative alterations of energy intake, energy expenditure or locomotion. Altogether, the present study supports the feasibility of biasing GHSR signaling to the benefit of the storage of fat by acting preferentially on nutrient partitioning.

The present study provides novel insights into the relationship between GHSR signaling and metabolic fuel preference. Indeed, *Ghsr*^M/M^ rats showed a higher RER than *Ghsr*^WT/WT^ littermates, indicating a shift in metabolic preference towards decreased use of fat. It is noteworthy that increased RER in *Ghsr*^M/M^ rats was essentially observed during light phase in the *ad libitum* condition (physiological fasting) and during early fasting and early refeeding. In the same time, circulating acyl ghrelin levels in *Ghsr*^M/M^ rats were similar or even lower than *Ghsr*^WT/WT^ littermates in the fed or fasted conditions, respectively. These observations are therefore in line with acute and chronic pharmacological studies showing that supraphysiological levels of acyl ghrelin elevates RER, indicating shifted metabolic fuel preference towards enhanced carbohydrate utilization, while energy expenditure and homecage activity are unaffected (Currie, et al. 2005; Theander-Carrillo et al. 2006; Tschop et al. 2000). Overall, based on the assets of the *Ghsr*^Q343X^ model enhancing ghrelin responsiveness (Chebani et al. 2016), the present data support a physiological role for GHSR signaling in regular diet conditions to promote fat storage and preservation by acting on nutrient partitioning, a mechanism that might be, at least in part, centrally mediated (Joly-Amado et al. 2012).

Metabolic characterization of *Ghsr*^M/M^ and *Ghsr*^WT/WT^ littermates explored herein supports a specific alteration of substrate balance (decreased fat oxidation) associated the *Ghsr*^Q343X^ mutation, rather than a modification of total energy intake or expenditure. Indeed, as measured in freely behaving conditions, *Ghsr*^M/M^ and *Ghsr*^WT/WT^ littermate rats showed similar chow intake, feeding patterns and energy expenditure. Although the latter did tend to decrease in *Ghsr*^M/M^ rats compared to *Ghsr*^WT/WT^ rats in situations of energy deficit, it did not differ in the fed condition, thus excluding a key role of GHSR signaling in energy expenditure control. Interestingly, young adult *Ghsr*^M/M^ rats showed increased body weight compared to their *Ghsr*^WT/WT^ littermates, but displayed equivalent proportions of fat and lean masses relatively to total body mass, as well as comparable fasting glycemia. In comparison, as exemplified in our prior study (Chebani et al. 2016), older *Ghsr*^M/M^ rats showed enhanced body weight and adiposity and decreased glucose tolerance, suggesting that the phenotype of these rats worsen with age, which is consistent with studies in *Ghsr*^-/-^ mice involving the GHSR in metabolic aging (Lin, et al. 2011; Ma, et al. 2011). Indeed, as they are aging, *Ghsr*^-/-^ mice fed on a standard diet display reduced obesity and improved insulin sensitivity compared to *Ghsr*^+/+^ littermates, a phenotype mirroring that of adult *Ghsr*^M/M^ rats. In contrast to the mechanism described in *Ghsr*^M/M^ rats (lower fat oxidation associated with development of adiposity), while RER of 1 year-old mice was similar amongst genotypes, it was increased in 2 year-old *Ghsr*^-/-^ mice compared to *Ghsr*^+/+^ littermates. It was suggested that increased RER in *Ghsr*^-/-^ mice could be related to their lean phenotype and improved insulin sensitivity (Lin et al. 2011). Altogether, the phenotype of *Ghsr*^M/M^ rats and *Ghsr*^-/-^ mice seems to implicate GHSR signaling in age-associated adiposity and insulin resistance. In addition, we speculate that enhanced acyl ghrelin sensitivity in *Ghsr*^Q343X^ rats illustrates how low fat oxidation may contribute to the occurrence of overweight (Galgani and Ravussin 2008).

The results obtained herein using pharmacological tools refine and further support previous observations on the mechanism-of-action of the *Ghsr*^Q343X^ mutation in rat. First, enhanced locomotor response to the GHSR agonist hexarelin in *Ghsr*^M/M^ rats may suggest increased dopaminergic responsiveness compared to *Ghsr*^WT/WT^ littermates. Former results using dose-responses of ghrelin or hexarelin already showed improved GH release and 4h chow intake, observations indicative of enhanced GHSR responsivity in the hypothalamus (Chebani et al. 2016). In sum, these experiments are consistent with the hypothesis that the *Ghsr*^Q343X^ mutation, that results in G protein biased signaling in response to agonist in cellular systems (Chebani et al. 2016), could recapitulate a gain-of-function mutation in the GHSR. Second, pharmacological challenges designed to probe dopaminergic circuits using a DAT blocker (amphetamine) or a DRD2 agonist (cabergoline) revealed no difference across *Ghsr*^M/M^ and *Ghsr*^WT/WT^ rats. These observations therefore suggest that the *Ghsr*^Q343X^ allele has no obvious direct or indirect effect regarding both of these pharmacological responses, involving enhanced extracellular dopamine tone (amphetamine) or DRD2-GHSR heterodimers (cabergoline) (Kern et al. 2012). Interestingly, rare human missense or nonsense *GHSR* variants segregating with short stature or GH deficiency, presumed to be loss-of-function mutations on the basis of their *in vitro* mechanism of action, were documented (Inoue, et al. 2011; Pantel, et al. 2009; Pantel, et al. 2006; Pugliese-Pires, et al. 2011), including the *GHSR*^A204E^ mutation that specifically alters constitutive activity *in vitro*. Just recently, a knock-in mice model expressing this mutation showed altered GH release, food intake and glycemic control (Torz, et al. 2020), therefore demonstrating that the *GHSR*^A204E^ disease-causing mutation is related to a partial impairment of GHSR functioning. This kind of genetic defect mirrors the mechanism of the present *Ghsr*^Q343X^ mutation in rat, documenting enhanced GHSR function both *in vitro* and *in vivo*. Altogether, *Ghsr* mutant models appear as very relevant tools to probe the significance of GHSR signaling *in vivo*, more especially since this constitutively active GPCR was recently established as the key target of both the agonist hormone ghrelin and LEAP2, an endogenous ligand with inverse agonist properties (M’Kadmi et al. 2019; Mani et al. 2019).

Using the *Ghsr*^Q343X^ model to probe food anticipatory activity, *Ghsr*^M/M^ and *Ghsr*^WT/WT^ rats were found to have similar running activity in anticipation of food, suggesting that the *Ghsr*^Q343X^ mutation does not impact the development nor expression of food anticipatory activity (FAA). Acyl ghrelin has been proposed to be a food-entrainable oscillator that participates in food anticipatory activity in rodents put on a restricted feeding schedule (Laermans, et al. 2015; LeSauter et al. 2009). Furthermore, when mice are put on a restricted feeding schedule, *Ghsr* knock-out results in attenuated FAA (Blum et al. 2009; Davis, et al. 2011; LeSauter et al. 2009) and reduced activation of several hypothalamic and midbrain nuclei prior to food access (Lamont, et al. 2012), altogether supporting that GHSR signaling plays a role in food anticipation. FAA was suggested to have a “go, no-go” property (LeSauter et al. 2009), therefore, while complete removal of the GHSR results in delayed onset of FAA, the presence of functional canonical acyl ghrelin-GHSR signaling in *Ghsr*^M/M^ rats, albeit enhanced (Chebani et al. 2016), might produce “go” decisions with probabilities similar to the wild type GHSR without affecting FAA onset. Of note, *Ghsr*^M/M^ rats were previously reported to have decreased FAA as assessed by total number of infrared bream breaks during the 2 hours preceding food access (MacKay, et al. 2016), a result that is not supported by present observations using two different paradigms (running wheels and actimetry cages). However, in this former study, FAA was not normalized to total 24h activity levels, and the observed decrease in FAA is likely to be the result of the general reduction in activity levels related to the metabolic phenotype of *Ghsr*^M/M^ rats, rather than a specific attenuation of food-oriented anticipatory activity.

Taking advantage of the high preference for sucrose of the Fawn hood strain (Tordoff et al. 2008) to probe spontaneous phenotypes in *ad libitum* fed *Ghsr*^M/M^ and *Ghsr*^WT/WT^ rats, the present study disclosed 1) similar consumption of sucrose in a two-bottle choice paradigm, suggesting that the hedonic perception of sucrose is not altered, and 2) similar levels of total nose-poke responses for sucrose in an instrumental task, supporting similar motivation to obtain food rewards. Thus far, pharmacological studies support that acyl ghrelin-GHSR signaling modulates the appetitive properties of palatable food. Indeed, peripherally administered acyl ghrelin promotes, whereas GHSR antagonist JMV2959 reduces, consumption and motivation to obtain palatable food in ad libitum-chow fed rodents (Landgren, et al. 2011; Perello, et al. 2010; Skibicka, et al. 2012), effects that are essentially reproduced by i.c.v., intra-VTA or intra-ventral hippocampus (vHPC) administration of acyl ghrelin or JMV2959 (Kanoski, et al. 2013; Skibicka et al. 2012; Skibicka, et al. 2011). However, the physiological significance of these effects is still unclear, as *Ghsr*^-/-^, *Ghrl*^-/-^ and *Goat*^-/-^ mouse models rarely showed results supporting altered hedonic feeding behaviors when animals are explored in the fed state (Davis, et al. 2012; Disse, et al. 2010; Lockie, et al. 2015). In the physiological setting of the present study, it is interesting to note that *Ghsr*^M/M^ rats, considered with enhanced response to endogenous ghrelin, did not produce an increase in the number of responses over *Ghsr*^WT/WT^ rats, but had a subtler effect on the performing speed of the animals, which was increased. This observation may suggest enhanced impulsivity to obtain food rewards. Interestingly, ghrelin injected i.c.v. or into the VTA of rats was indeed recently shown to increase impulsive behavior to obtain palatable food (Anderberg, et al. 2016). Overall, the observation supporting a possible effect of the *Ghsr*^Q343X^ mutation with qualitative rather than a quantitative modulation of spontaneous operant responding for sucrose, at least in female rats, is of particular interest and needs to be clarified further, keeping in mind possible confounding factor such as the current rat strain as well as the metabolic phenotype of aged *Ghsr*^M/M^ rats.

Altogether, the present study strengthens the observations that *Ghsr*^M/M^ rats may specifically show enhanced responsivity to GHSR agonist *in vivo*. In the fed or fasted conditions, young adult *Ghsr*^M/M^ rats did not show obvious feeding or locomotor alterations but showed shifted fuel preference towards carbohydrates, therefore providing a possible mechanism to the enhanced fat accumulation documented in these rats later with age. Finally, these data also support the feasibility of tricking GHSR signaling to the benefit of fat storage by acting preferentially on nutrient partitioning.

## Author contribution statement

C.M., P.Z., R.G.P.D, J.P. conceptualized the research, C.M., P.Z., R.G.P.D., R.H., Y.C., J.P. conducted the experiments, C.M., P.Z., R.G.P.D., J.P. analyzed the data, G.L.P, F.N., S.L., J.P. supervised the experiments, C.M. and J.P. wrote the initial draft of the manuscript. All authors edited and revised the manuscript.

## Acknowledgements

The authors are grateful to the PhysGen program for providing the *Ghsr* FHH rats (Dr. Howard Jacob, Medical College of Wisconsin), to the animal core and PhenoBrain facilities of the Institute of Psychiatry and Neuroscience of Paris (INSERM UMR S-1266 | Université de Paris), the animal core facility of BioMedTech (INSERM US36 | CNRS UMS2009 | Université de Paris). The authors are also thankful to Corinne Canestrelli, Claire Dovergne, Racha Fayad and Adèle Guillard for technical support, to Gabriel Montaldo for help with activity wheel recording. We also acknowledge the technical platform metabolism of the Unit “Biologie Fonctionnelle et Adaptative” (University de Paris, BFA UMR CNRS 8251) for metabolic analysis and the animal core facility “Buffon” of the University Paris Diderot Paris 7/Institut Jacques Monod, Paris for animal housing.

---

**Supplementary Figure 1:**
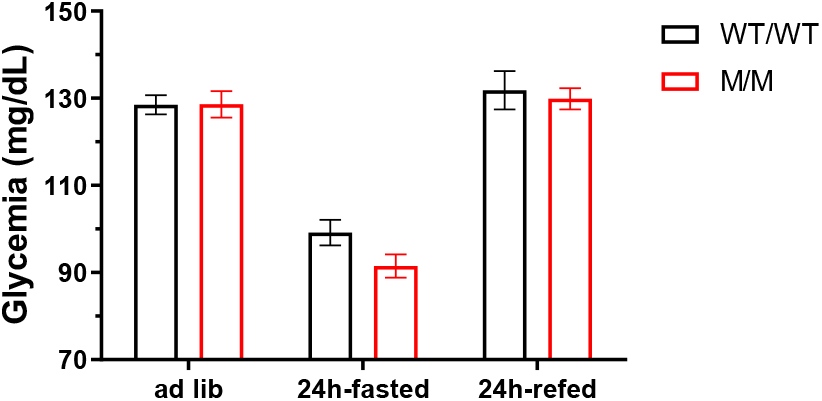
Glycemia fluctuations across feeding conditions. *Ghsr*^M/M^ and *Ghsr*^WT/WT^ male rats at 15 weeks of age (n=6-8/genotype). Data represent mean ± SEM.

**Supplementary Figure 2:**
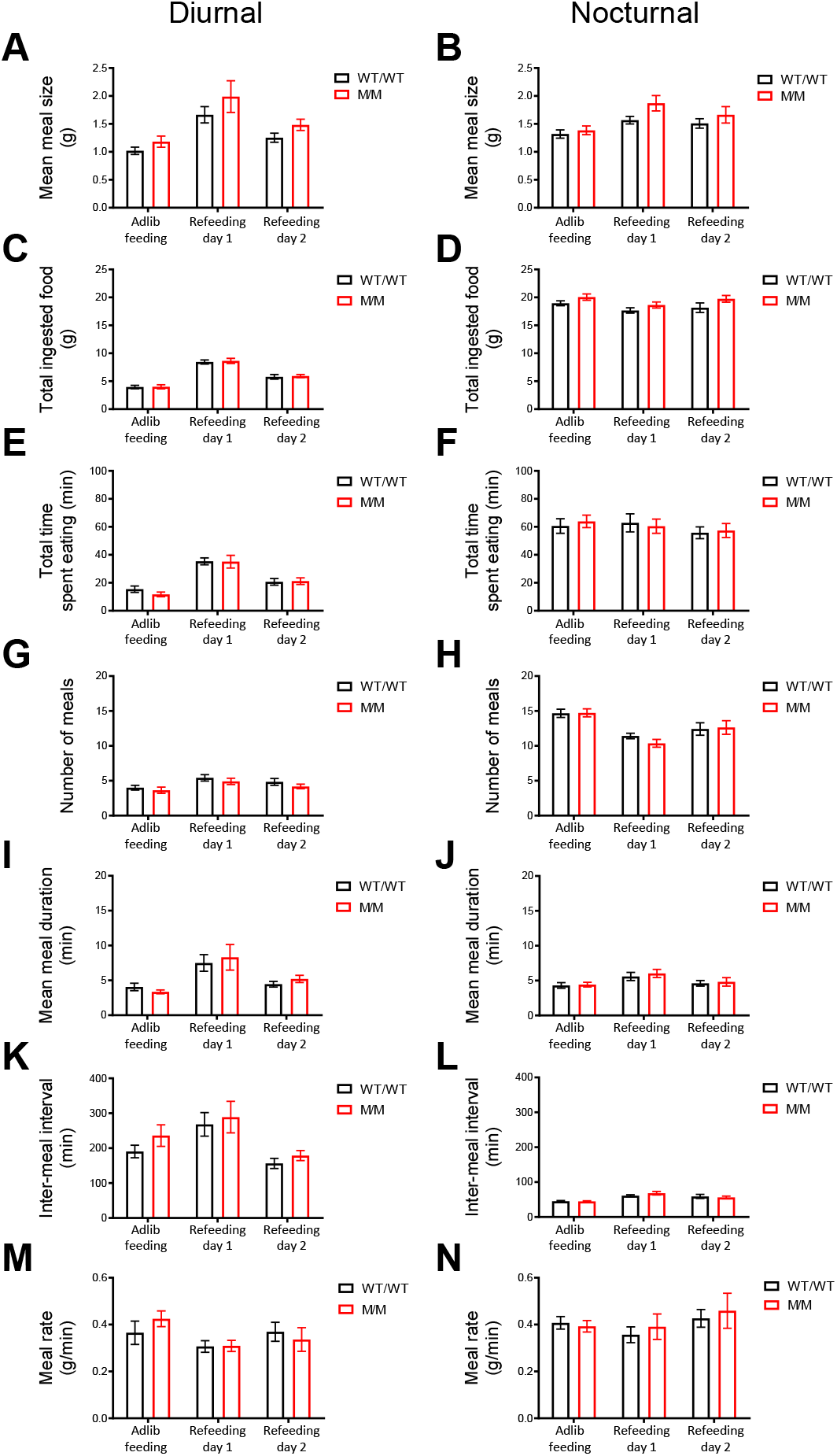
Diurnal and nocturnal meal patterns of *Ghsr*^M/M^ and *Ghsr*^WT/WT^ rats during ad libitum-feeding and refeeding. (A-B) Mean meal size during light (A) and dark (B) phases. (C-D) Total chow ingested during light (C) and dark (D) phases. (E-F) Total time spent eating during light (E) and dark (F) phases. (G-H) Number of meals during light (G) and dark (H) phases. (I-J) Mean meal duration in light (I) and dark (J) phases. (K-L) Inter-meal interval during light (K) and dark (L) phases. (M-N) Meal ingestion rate in light (M) and dark (N) phases (n=12/genotype). Data represent mean ± SEM.

**Supplementary Figure 3:**
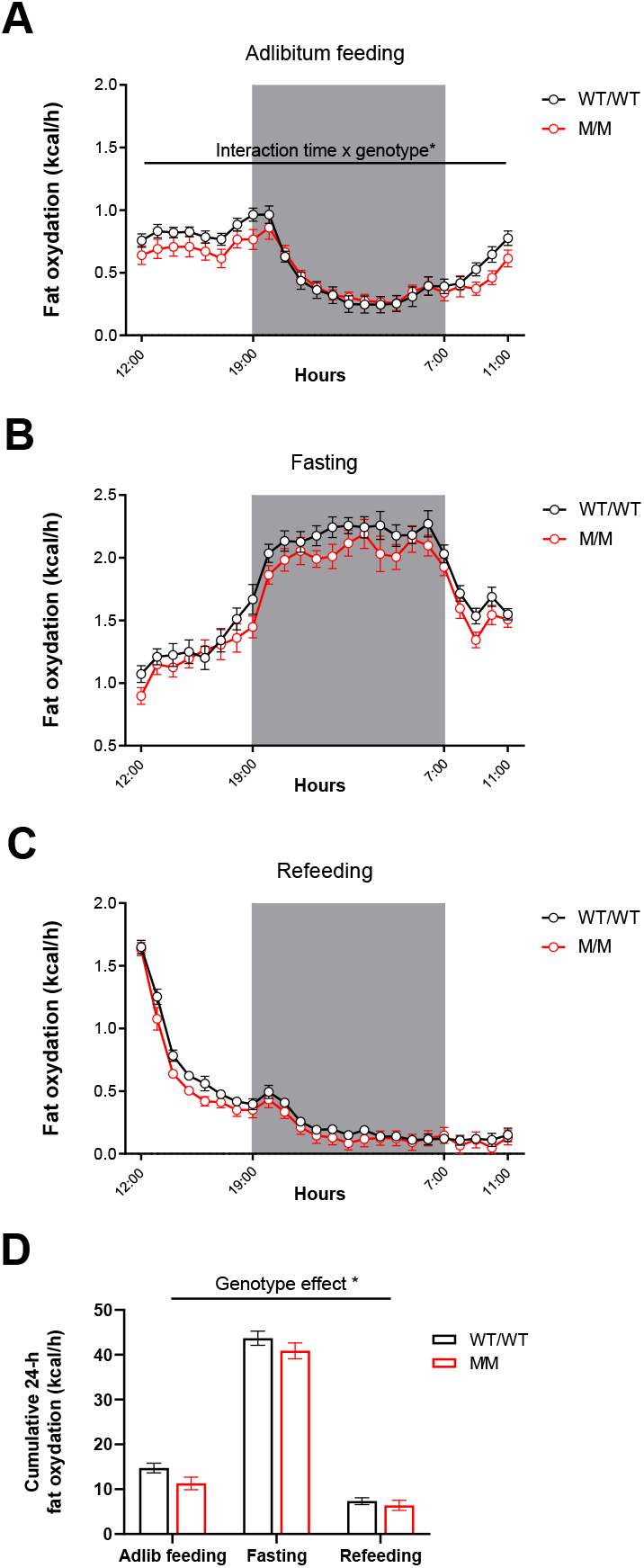
Decreased fat oxydation in *Ghsr*^M/M^ rats across feeding conditions. (A-C) Daily pattern of fat oxydation during adlibitum feeding (A), 24h fasting (B) and 24h refeeding (C). Data represent mean ± SEM. (D) 24-h cumulative fat oxydation (n=12/genotype). Data were analyzed by 2-way ANOVA. * p<0.05.

## References

Abizaid A, Liu ZW, Andrews ZB, Shanabrough M, Borok E, Elsworth JD, Roth RH, Sleeman MW, Picciotto MR, Tschop MH, et al. 2006 Ghrelin modulates the activity and synaptic input organization of midbrain dopamine neurons while promoting appetite. Journal of Clinical Investigation 116 3229–3239.

Al-Massadi O, Muller T, Tschop M, Dieguez C & Nogueiras R 2018 Ghrelin and LEAP-2: Rivals in Energy Metabolism. Trends in Pharmacologicol Sciences 39 685–694.

Anderberg RH, Hansson C, Fenander M, Richard JE, Dickson SL, Nissbrandt H, Bergquist F & Skibicka KP 2016 The Stomach-Derived Hormone Ghrelin Increases Impulsive Behavior. Neuropsychopharmacology 41 1199–1209.

Andrews ZB 2019 The next big LEAP2 understanding ghrelin function. Journal of Clinical Investigation 129 3542–3544.

Blum ID, Patterson Z, Khazall R, Lamont EW, Sleeman MW, Horvath TL & Abizaid A 2009 Reduced anticipatory locomotor responses to scheduled meals in ghrelin receptor deficient mice. Neuroscience 164 351–359.

Chebani Y, Marion C, Zizzari P, Chettab K, Pastor M, Korostelev M, Geny D, Epelbaum J, Tolle V, Morisset-Lopez S, et al. 2016 Enhanced responsiveness of *Ghsr*^Q343X^ rats to ghrelin results in enhanced adiposity without increased appetite. Science Signaling 9 ra39.

Currie PJ, Mirza A, Fuld R, Park D & Vasselli JR 2005 Ghrelin is an orexigenic and metabolic signaling peptide in the arcuate and paraventricular nuclei. American Journal of Physiology: Regulatory, Integrative and Comparative Physiology 289 R353–R358.

Damian M, Marie J, Leyris JP, Fehrentz JA, Verdie P, Martinez J, Baneres JL & Mary S 2012 High constitutive activity is an intrinsic feature of ghrelin receptor protein: a study with a functional monomeric GHS-R1a receptor reconstituted in lipid discs. Journal of Biological Chemistry 287 3630–3641.

Davis JF, Choi DL, Clegg DJ & Benoit SC 2011 Signaling through the ghrelin receptor modulates hippocampal function and meal anticipation in mice. Physiology & Behavior 103 39–43.

Davis JF, Perello M, Choi DL, Magrisso IJ, Kirchner H, Pfluger PT, Tschoep M, Zigman JM & Benoit SC 2012 GOAT induced ghrelin acylation regulates hedonic feeding. Hormones and Behavior 62 598–604.

Disse E, Bussier AL, Veyrat-Durebex C, Deblon N, Pfluger PT, Tschop MH, Laville M & Rohner-Jeanrenaud F 2010 Peripheral ghrelin enhances sweet taste food consumption and preference, regardless of its caloric content. Physiology & Behavior 101 277–281.

Galgani J & Ravussin E 2008 Energy metabolism, fuel selection and body weight regulation. International Journal of Obesity 32 Suppl 7 S109–119.

Ge X, Yang H, Bednarek MA, Galon-Tilleman H, Chen P, Chen M, Lichtman JS, Wang Y, Dalmas O, Yin Y, et al. 2018 LEAP2 Is an Endogenous Antagonist of the Ghrelin Receptor. Cell Metabolism 27 461–469 e466.

Holst B, Cygankiewicz A, Jensen TH, Ankersen M & Schwartz TW 2003 High constitutive signaling of the ghrelin receptor--identification of a potent inverse agonist. Molecular Endocrinology 17 2201–2210.

Howard AD, Feighner SD, Cully DF, Arena JP, Liberator PA, Rosenblum CI, Hamelin M, Hreniuk DL, Palyha OC, Anderson J, et al. 1996 A receptor in pituitary and hypothalamus that functions in growth hormone release. Science 273 974–977.

Inoue H, Kangawa N, Kinouchi A, Sakamoto Y, Kimura C, Horikawa R, Shigematsu Y, Itakura M, Ogata T & Fujieda K 2011 Identification and functional analysis of novel human growth hormone secretagogue receptor (GHSR) gene mutations in Japanese subjects with short stature. Journal of Clinical Endocrinology & Metabolism 96 E373–378.

Jerlhag E, Egecioglu E, Dickson SL, Douhan A, Svensson L & Engel JA 2007 Ghrelin administration into tegmental areas stimulates locomotor activity and increases extracellular concentration of dopamine in the nucleus accumbens. Addiction Biology 12 6–16.

Joly-Amado A, Denis RG, Castel J, Lacombe A, Cansell C, Rouch C, Kassis N, Dairou J, Cani PD, Ventura-Clapier R, et al. 2012 Hypothalamic AgRP-neurons control peripheral substrate utilization and nutrient partitioning. EMBO Journal 31 4276–4288.

Kanoski SE, Fortin SM, Ricks KM & Grill HJ 2013 Ghrelin signaling in the ventral hippocampus stimulates learned and motivational aspects of feeding via PI3K-Akt signaling. Biological Psychiatry 73 915–923.

Kern A, Albarran-Zeckler R, Walsh HE & Smith RG 2012 Apo-ghrelin receptor forms heteromers with DRD2 in hypothalamic neurons and is essential for anorexigenic effects of DRD2 agonism. Neuron 73 317–332.

Kojima M, Hosoda H, Date Y, Nakazato M, Matsuo H & Kangawa K 1999 Ghrelin is a growth-hormone-releasing acylated peptide from stomach. Nature 402 656–660.

Laermans J, Vancleef L, Tack J & Depoortere I 2015 Role of the clock gene Bmal1 and the gastric ghrelin-secreting cell in the circadian regulation of the ghrelin-GOAT system. Scientific Reports 5 16748.

Lamont EW, Patterson Z, Rodrigues T, Vallejos O, Blum ID & Abizaid A 2012 Ghrelin-deficient mice have fewer orexin cells and reduced cFOS expression in the mesolimbic dopamine pathway under a restricted feeding paradigm. Neuroscience 218 12–19.

Landgren S, Simms JA, Thelle DS, Strandhagen E, Bartlett SE, Engel JA & Jerlhag E 2011 The ghrelin signalling system is involved in the consumption of sweets. PLoS One 6 e18170.

LeSauter J, Hoque N, Weintraub M, Pfaff DW & Silver R 2009 Stomach ghrelin-secreting cells as food-entrainable circadian clocks. Proceedings of the National Academy of Sciences of the United States of America 106 13582–13587.

Lin L, Saha PK, Ma X, Henshaw IO, Shao L, Chang BH, Buras ED, Tong Q, Chan L, McGuinness OP, et al. 2011 Ablation of ghrelin receptor reduces adiposity and improves insulin sensitivity during aging by regulating fat metabolism in white and brown adipose tissues. Aging Cell 10 996–1010.

Lockie SH, Dinan T, Lawrence AJ, Spencer SJ & Andrews ZB 2015 Diet-induced obesity causes ghrelin resistance in reward processing tasks. Psychoneuroendocrinology 62 114–120.

M’Kadmi C, Cabral A, Barrile F, Giribaldi J, Cantel S, Damian M, Mary S, Denoyelle S, Dutertre S, Peraldi-Roux S, et al. 2019 N-Terminal Liver-Expressed Antimicrobial Peptide 2 (LEAP2) Region Exhibits Inverse Agonist Activity toward the Ghrelin Receptor. Journal of Medicinal Chemistry 62 965–973.

Ma X, Lin L, Qin G, Lu X, Fiorotto M, Dixit VD & Sun Y 2011 Ablations of ghrelin and ghrelin receptor exhibit differential metabolic phenotypes and thermogenic capacity during aging. PLoS One 6 e16391.

MacKay H, Charbonneau VR, St-Onge V, Murray E, Watts A, Wellman MK & Abizaid A 2016 Rats with a truncated ghrelin receptor (GHSR) do not respond to ghrelin, and show reduced intake of palatable, high-calorie food. Physiology & Behavior 163 88–96.

Mani BK & Zigman JM 2017 Ghrelin as a Survival Hormone. Trends in Endocrinology and Metabolism 28 843–854.

Mani BK, Puzziferri N, He Z, Rodriguez JA, Osborne-Lawrence S, Metzger NP, Chhina N, Gaylinn B, Thorner MO, Thomas EL, et al. 2019 LEAP2 changes with body mass and food intake in humans and mice. Journal of Clinical Investigation 129 3909–3923.

Muller TD, Nogueiras R, Andermann ML, Andrews ZB, Anker SD, Argente J, Batterham RL, Benoit SC, Bowers CY, Broglio F, et al. 2015 Ghrelin. Molecular Metabolism 4 437–460.

Nakazato M, Murakami N, Date Y, Kojima M, Matsuo H, Kangawa K & Matsukura S 2001 A role for ghrelin in the central regulation of feeding. Nature 409 194–198.

Naleid AM, Grace MK, Cummings DE & Levine AS 2005 Ghrelin induces feeding in the mesolimbic reward pathway between the ventral tegmental area and the nucleus accumbens. Peptides 26 2274–2279.

Pantel J, Legendre M, Nivot S, Morisset S, Vie-Luton MP, le Bouc Y, Epelbaum J & Amselem S 2009 Recessive isolated growth hormone deficiency and mutations in the ghrelin receptor. Journal of Clinical Endocrinology & Metabolism 94 4334–4341.

Pantel J, Legendre M, Cabrol S, Hilal L, Hajaji Y, Morisset S, Nivot S, Vie-Luton MP, Grouselle D, de Kerdanet M, et al. 2006 Loss of constitutive activity of the growth hormone secretagogue receptor in familial short stature. Journal of Clinical Investigation 116 760–768.

Perello M, Cabral A, Cornejo MP, De Francesco PN, Fernandez G & Uriarte M 2019 Brain accessibility delineates the central effects of circulating ghrelin. Journal of Neuroendocrinology 31 e12677.

Perello M, Sakata I, Birnbaum S, Chuang JC, Osborne-Lawrence S, Rovinsky SA, Woloszyn J, Yanagisawa M, Lutter M & Zigman JM 2010 Ghrelin increases the rewarding value of high-fat diet in an orexin-dependent manner. Biological Psychiatry 67 880–886.

Pugliese-Pires P, Fortin JP, Arthur T, Latronico AC, de Mendonca BB, Villares SM, Arnhold IJ, Kopin AS & Jorge AA 2011 Novel Inactivating Mutations in the Growth Hormone Secretatogue Receptor Gene (GHSR) in Patients with Constitutional Delay of Growth and Puberty. European journal of Endocrinology 165 233–41.

Riachi M, Himms-Hagen J & Harper ME 2004 Percent relative cumulative frequency analysis in indirect calorimetry: application to studies of transgenic mice. Canadian Journal of Physiology and Pharmacology 82 1075–1083.

Skibicka KP, Hansson C, Egecioglu E & Dickson SL 2012 Role of ghrelin in food reward: impact of ghrelin on sucrose self-administration and mesolimbic dopamine and acetylcholine receptor gene expression. Addiction Biology 17 95–107.

Skibicka KP, Hansson C, Alvarez-Crespo M, Friberg PA & Dickson SL 2011 Ghrelin directly targets the ventral tegmental area to increase food motivation. Neuroscience 180 129–137.

Theander-Carrillo C, Wiedmer P, Cettour-Rose P, Nogueiras R, Perez-Tilve D, Pfluger P, Castaneda TR, Muzzin P, Schurmann A, Szanto I, et al. 2006 Ghrelin action in the brain controls adipocyte metabolism. Journal of Clinical Investigation 116 1983–1993.

Tordoff MG, Alarcon LK & Lawler MP 2008 Preferences of 14 rat strains for 17 taste compounds. Physiology & Behavior 95 308–332.

Torz LJ, Osborne-Lawrence S, Rodriguez J, He Z, Cornejo MP, Mustafa ER, Jin C, Petersen N, Hedegaard MA, Nybo M, et al. 2020 Metabolic insights from a GHSR-A203E mutant mouse model. Molecular Metabolism 39 101004.

Tschop M, Smiley DL & Heiman ML 2000 Ghrelin induces adiposity in rodents. Nature 407 908–913.

Wang Q, Liu C, Uchida A, Chuang JC, Walker A, Liu T, Osborne-Lawrence S, Mason BL, Mosher C, Berglund ED, et al. 2014 Arcuate AgRP neurons mediate orexigenic and glucoregulatory actions of ghrelin. Molecular Metabolism 3 64–72.

Zigman JM, Jones JE, Lee CE, Saper CB & Elmquist JK 2006 Expression of ghrelin receptor mRNA in the rat and the mouse brain. Journal of Comparative Neurology 494 528–548.

